# Effect of altered production and storage of dopamine on development and behavior in *C. elegans*

**DOI:** 10.1101/2023.10.07.561350

**Authors:** Irene Lee, Ava C Knickerbocker, Charlotte Rose Depew, Elizabeth Martin, Jocelyn Dicent, Gary W Miller, Meghan L Bucher

**Author notes:** indicates equal contribution (co-first authorship) (Irene Lee, Ava C Knickerbocker).

## Abstract

The nematode, *Caenorhabditis elegans*, is an advantageous model for studying developmental toxicology due to its homology to humans and well-defined developmental stages. Similarly to humans, *C. elegans* utilize dopamine as a neurotransmitter to regulate motor behavior. We have previously reported behavioral deficits in a genetic model of *C. elegans* (OK411) that lack the neurotransmitter transporter necessary for packaging dopamine into synaptic vesicles. Anecdotally, we observed these *C. elegans* appeared to have a smaller body size, which is supported by prior studies that observed a larger body size in *C. elegans* that lack the enzyme that catalyzes dopamine synthesis, suggesting a complex regulatory system in which dopamine mediates body size in *C. elegans*. However, the question of whether body size abnormalities apparent in *C. elegans* with disruptions to their dopamine system are developmental or purely based on body size remains unanswered. Here, we present data characterizing the effect of gene mutations in dopamine-related proteins on body size, development, and behavior. We analyzed *C. elegans* that lack the ability to sequester dopamine (OK411), that overproduce dopamine (UA57), and a novel strain (MBIA) generated through crossing OK411 and UA57, which lacks the ability to sequester dopamine into vesicles and additionally endogenously overproduces dopamine. This novel strain was generated to address the hypothesis that an endogenous increase in production of dopamine can rescue deficits caused by a lack of vesicular dopamine sequestration. Compared to wild type, OK411 have shorter body lengths and behavioral deficits in early life stages. In contrast, the MBIA strain have similar body lengths to wild-type by early adulthood and display similar behavior to wild-type by early adulthood. Our data suggests that endogenously increasing the production of dopamine is able to mitigate deficits in *C. elegans* lacking the ability to package dopamine into synaptic vesicles. These results provide evidence that the dopamine system impacts development, growth, and reproduction in *C. elegans*.

## Introduction

The model organism *Caenorhabditis Elegans (C. elegans)* has significant utility for developmental toxicology research due to their small size, transparent body, defined developmental staging, and eutelic nature.^1,2^ In comparison to the complex dopamine systems consisting of thousands of dopaminergic neurons in the human brain, *C. elegans* have a simple dopaminergic system consisting of eight neurons - six in the head and two in the body.^3^ Despite the difference in complexity and number of neurons, dopamine regulates similar behaviors in *C. elegans* as in humans including locomotion, food response, enhancement of odor avoidance, mating behavior, plasticity of mechanosensory response, and behavioral choice.^4-9^ In humans, dysregulation of dopamine homeostasis has been linked to a number of diseases and disorders of the dopamine system including attention deficit hyperactivity disorder, substance use disorder, and Parkinson’s disease.

In humans, dopamine synthesis occurs when tyrosine is converted to levodopa (L-DOPA) by tyrosine hydroxylase (TH) and L-DOPA is decarboxylated to dopamine by aromatic amino acid decarboxylase.^10,11^ Following synthesis, dopamine is sequestered into synaptic vesicles by vesicular monoamine transporter 2 (VMAT2), which is a necessary process for proper neurotransmission and to limit the amount of free dopamine in the cytosol that has the potential to be neurotoxic.^12^ *C. elegans* contain homologs for human TH and VMAT2 encoded by the *cat-2* and *cat-1* genes, respectively.^13,14^ The VMAT2 homolog in *C. elegans, cat-1*, was reported to be 47% identical to human VMAT1 and 49% identical to human VMAT2, demonstrating uptake of dopamine and serotonin.^14-16^ Furthermore, the uptake of these substrates was inhibited by known VMAT inhibitors tetrabenazine and reserpine demonstrating a similar inhibitor profile to that of mammalian VMAT2.^14,15^ Due to the high degree of conserved biology between *C. elegans* and humans, and the genetic tractability of *C. elegans*, transgenic strains with gene mutations in proteins involved in dopamine homeostasis can be easily generated to investigate human conditions.

In *C. elegans, cat-1* null animals demonstrate minor egg laying deficits and decreased mating performance in males.^14,17^ These animals also display age-dependent degeneration and increased susceptibility to the dopaminergic neurotoxicant and VMAT2 substrate MPP^+^.^18^ These findings substantiate previous work from rodent models demonstrating that manipulating VMAT2 expression can alter the amount of dopamine in the cytosol, mediate neuronal vulnerability to degeneration particularly in cases of toxicant exposures, and cause behavioral effects.^19-27^ In Parkinson’s disease, there is significant evidence of dysfunctional VMAT2 expression and activity, suggesting a pathogenic role of dysregulated dopamine sequestration in disease onset and/or progression.^28-30^ The motor deficits resulting from deficient dopamine transmission in Parkinson’s disease are treated by supplementing with the precursor to dopamine, L-DOPA.^31,32^ In *C. elegans, cat-2* overexpression is used as an endogenous model of L-DOPA exposure to replicate an increase in dopamine production; however, these animals show age-dependent dopaminergic neurodegeneration.^33,34^

Taken together, previous studies have found that proper dopamine synthesis and sequestration is essential to maintaining neuronal health in C. elegans, and dysregulation of dopamine homeostasis can affect many behaviors in *C. elegans* including growth and reproduction.. In this study, we sought to determine whether endogenously increasing dopamine production through the overexpression of *cat*-*2* could rescue the deficits identified in *cat-1* null *C. elegans*. To do this, we bred two parent *C. elegans* strains OK411 (*cat-1*^*null*^) and UA57 (*cat-2*^*OE*^) to generate a novel strain called MBIA (*cat-1*^*null*^*::cat-2*^*OE*^). Here, we present a thorough characterization of the MBIA (*cat-1*^*null*^*::cat-2*^*OE*^) strain investigating body size, reproduction, neuron health, and dopamine-mediated behaviors.

## Methods

### *C*. *elegans* husbandry

*C. elegans* were maintained on agar plates with normal growth medium (NGM) at 20ºC with the OP50 strain of *e. coli* as food.

### Synchronization of population

Synchronization of *C. elegans* populations was performed through the following protocol. Gravid adult *C. elegans* were collected from agar plates using M9 buffer (22mM KH_2_PO_4_, 42mM Na_2_HPO_4_, 85mM NaCl, 1mM MgSO_4_ prepared in sterile water) and collected into 1.5mL Eppendorf tubes. Tubes were centrifuged at 11,000 rpm for 1 min, supernatant was removed, and 1mL of bleach solution (250mM KOH, 0.5% hypochlorite prepared in sterile water) was added to each tube. Tubes containing bleach and *C. elegans* were rested on the bench on their side with periodic vortexing until most of the bodies disintegrated. Tubes were centrifuged at 11,000 rpm for 1 min, bleach supernatant was discarded, and eggs underwent two washes to remove any residual bleach solution by resuspending in 1mL of M9 and centrifuging at 11,000 prm for 1 min. Following the last wash, supernatant was removed, and eggs were resuspended in 100μL of M9 before transferring to 50mL conical tubes in a final volume of 2mL M9 solution. Conical tubes were kept on a shaking incubator at 120 rpm and 20ºC overnight to let eggs hatch to the Larval 1 stage (L1). L1 *C. elegans* were placed on 6cm agar growth plates (normal growth medium with live *e. coli* OP50) and kept upside down in a 20°C incubator. For L4 analysis, *C. elegans* were collected 48 hours after L1 plating. Day One Adult *C. elegans* were analyzed 72 hours after L1 plating. Day Two Adult *C. elegans* were analyzed 96 hours after L1 plating. Day Three Adult *C. elegans* were analyzed 120 hours after L1 plating.

### Genotype confirmation by PCR

For the confirmation of genotype of MBIA strain, PCR was performed by collecting ∼5 gravid adult *C. elegans* for analysis with Sigma Extract-N-AMP™ Tissue PCR Kit (XNAT2) and Apex Taq Red 2X Master Mix. Briefly, worm samples were placed in lysis buffer and incubated in thermocycler at 55 ºC for 10m and 95ºC for 3m. Following incubation, neutralization solution was added to each tube to halt reaction. A primer set was used to confirm gene deletion of *cat-1* with a 701 base pair product in wild-type, and a 272 base pair product in *C. elegans* with *cat-1* null. Forward-3 primer: CCGCGCAAATGAATGACCTA. Reverse-3 primer: GAACCGGAAGGTTCAAGCAT. Worm lysate was mixed with Apex Taq Red 2X Master Mix, primers, and water, and underwent thermocycling of: 94ºC 2min, 94ºC (30sec 35x), 55.6ºC (30sec 35x), 72ºC (1min 35x), 72ºC (5min), 4ºC (hold). PCR product was run with loading buffer on pre-cast 2% gels (ThermoFisher) using the E-gel Power Snap Electrophoresis System (ThermoFisher).

### Microscopy analysis

*C. elegans* were imaged on microscope slides with agar pads (3% pure agarose in M9). Agar mixture was brought to a boil before allowing to cool to 58°C. Cooled agar was pipetted onto a glass microscopy slide and a second slide was placed on top to flatten the agar into an even surface. After the agar cooled completely and solidified, the top microscope slide was removed and 15-30 *C. elegans* were placed on the agar pad in 5mM Levamisole before applying a coverslip. Slides were imaged on an EVOS_*fl*_ or APEX100 microscope using Brightfield or GFP filter cubes at 4x, 10x, and 20x objectives.

#### Body Size

*C. elegans* on microscope slides were imaged on an EVOS_fl_ microscope at 4x using brightfield for body size analysis. Images were analyzed using ImageJ and FIJI by using the segmented line tool and drawing a line through the midline of the *C. elegans* from head to tail. The length of the segmented line was recorded in microns as a measurement of the body length of *C. elegans*.

#### Neuron Size

*C. elegans* on microscope slides were imaged on an EVOS_fl_ microscope at 20x using GFP filter at consistent light intensity and exposure. The optimal z-plane was determined by finding the plane where the most neurons in the head were in focus for imaging. ImageJ and FIJI were used to create and apply a macro to consistently analyze neuron area size across different worm ages and strains. Each image of a worm’s neurons was converted from GFP to grayscale (Image→Type→8-bit). Thresholding values to allow measurement of neuron size from the grayscale images were applied (Image→Adjust→Threshold [Select Default, Select B&W, Upper Value = 56, Lower Value = 200]→Apply). Upper Value and Lower Value values were determined by testing various thresholding sizes in random images, before determining that Upper Value = 56 and Lower Value = 200. Once the thresholding values were set, the ImageJ measured the area of the neurons in each image (Analyze→Analyze Particles (Size: 0.50-Infinity; Display Results; Show: Outlines)→OK). To record the macro, an image was opened in ImageJ then the steps that would appear in the Macro were recorded (Plugins→Macros→Record→Image→Adjust→Thres hold→Analyze Particles →Create→Save as a download through R-studios). The Macro was then applied to each image (Plugins →Macros→Run→Select Macro from files). After applying the Macro to each image, two additional windows were created in addition to the image of the worm’s neurons. The first tab displayed outlines of all noticeable areas that were measured, with a number in the center corresponding to what Area Size/Mean/Min/Max it possessed on the second tab.

The second tab was created and displayed the Area Size, Mean, Min, and Max values for each noticeable area. To exclude the areas of anything other than the neurons (e.g., scale bar, tail neurons of neighboring worm), the numbers that corresponded to the area of the neurons were manually confirmed. The areas of the confirmed neurons were copied and pasted onto a document recording all the data and manual exclusion of any objects less than 10 was performed.

### Behavior

For behavioral analysis, 4-6 *C. elegans* of a specific genotype and developmental stage were placed in 60μL of M9 solution added to microscope slides with hydrophobic circles. A stereo microscope with a camera attachment was used to analyze the behavior of *C. elegans* . The microscope was focused such that a well-defined image could be captured, and videos of *C. elegans* swimming in M9 were collected for 30,000ms at 18fps and saved as JPEG images. 4-6 replicates were performed per strain. Behavioral analysis was performed via ImageJ. A folder of images was imported as a sequence (File→Import→Image Sequence). Once images were processed, the zoom feature was used to zoom in on a worm at a setting of 300x. To measure the worm’s length, the full length of the worm was outlined using the segmented line feature and its subsequent length recorded (analyze→measure). To perform curling analysis, the image sequence was played through and the number of times an individual worm was in a curled position was counted. We defined curling as the position in which a worm’s tail was directly crossing its body.^35^ To perform thrashing analysis, we ensured that animation options were set to 18fps so that the image sequence played a 30 second video. A stopwatch starting at 0 sec. was prepared prior to starting the video. The percentage of time a worm was thrashing over a 30s time period was measured. We defined thrashing as the motion of a worm aggressively moving side to side.^35^ The length, curling, and thrashing analysis was repeated for each individual worm.

### Fecundity assay

Individual *C. elegans* were placed in one well of a clear U-shaped 96-well plate filled with 100μL of a 20% hypochlorite solution (in M9) per well. Plate was observed in brightfield at 4x and 10x on EVOS_*fl*_ microscope. Once worm body dissolved and eggs were released, eggs were counted and recorded as a measure of eggs *in utero*.

### COPAS

*C. elegans* at the L4 and Day One Adult developmental stages were analyzed using the Complex Object Parametric Analyzer and Sorter Flow Pilot-250 (COPAS FP-250) large particle flow cytometer (Union Biometrica, Massachusetts). *C. elegans* were collected from agar growth plates using 5mM Levamisole to anesthetize the animals. Processing through the COPAS recorded each object’s extinction and time of flight, which was exported in .csv files, which were analyzed using RStudio. Raw data files from each strain were stored in separate variables and then combined into one .csv file per experiment containing data for all strains using the “rbind” function. To exclude objects that were not *C. elegans* (e.g., contaminants, pieces of agar, dust particles, etc.), the Extinction measurements were filtered to restrict objects smaller than 250 and greater than 4000. A final .csv file with the cleaned data was generated and imported into Graph Pad Prism 10 for statistical analysis.

### Statistics

All experiments and analyses were performed blinded to genotype. All data were analyzed using GraphPad Prism 10 statistical software package (San Diego, California). Two-way ANOVA followed by Tukey’s multiple comparisons test was performed in each case where more than two genotypes were being compared across multiple developmental stages. For neuron area analysis, Two-way ANOVA with Šídák’s multiple comparisons test was performed. For COPAS analysis, One-way ANOVA followed by Tukey’s multiple comparisons test was performed.

## Results

### Creation of novel C. elegans strain “MBIA” overexpressing cat-2 with cat-1 null

The novel *C. elegans* strain “MBIA” was generated by crossing the two previously existing strains UA57 (*cat-2* overexpressing; *cat-2*^*OE*^) and OK411 (*cat-1* null; *cat-1*^*n*ull^) (Figure 1A). In addition to overexpressing *cat-2*, an enzyme involved in dopamine synthesis, UA57 *C. elegans* possess fluorescent dopaminergic neurons expressing green fluorescent protein (GFP). Male UA57 *C. elegans* were generated by heatshock protocol. Briefly, 10-15 L4 hermaphrodite UA57 *C. elegans* were placed on an agar plate with OP50 food in a 30°C incubator for 4h. The plate was transferred to standard incubation environment of 20°C and *C. elegans* were allowed to reproduce. Male UA57 *C. elegans* were identified by phenotype (e.g., fin-shaped tail) and placed on an agar plate with OP50 with hermaphrodite *C. elegans* to generate a consistent source of male UA57 offspring. Male UA57 *C. elegans* were placed on an agar plate with hermaphrodite OK411 *C. elegans* to promote breeding to generate novel MBIA strain containing both the UA57 and OK411 genotypes. Individual first-generation offspring (F1) from the UA57 x OK411 cross with fluorescent dopaminergic neurons indicating the presence of UA57 genotype were selected and placed on agar plates with OP50. These individual F1 *C. elegans* were allowed to hermaphroditically reproduce generating second-generation offspring (F2). Based on principles of Mendelian inheritance, F2 offspring were hypothesized to be present in a 1:2:1 ratio of homozygous for wild-type *cat-1* alleles, heterozygous for *cat-1* alleles, and homozygous for *cat-1* null. Thus, individual F2 offspring with fluorescent dopaminergic neurons indicating the presence of UA57 genotype were selected and placed on agar plates with OP50 and allowed to reproduce. Once the individual F2 worm had reproduced, the parent F2 worm was collected for PCR analysis to confirm the genotype of the strain. Fluorescent dopaminergic neurons confirmed the presence of UA57 phenotype, and through PCR analysis, the *cat-1* null genotype was identified by shift in genome size (Figure 1B).

**Figure 1.**
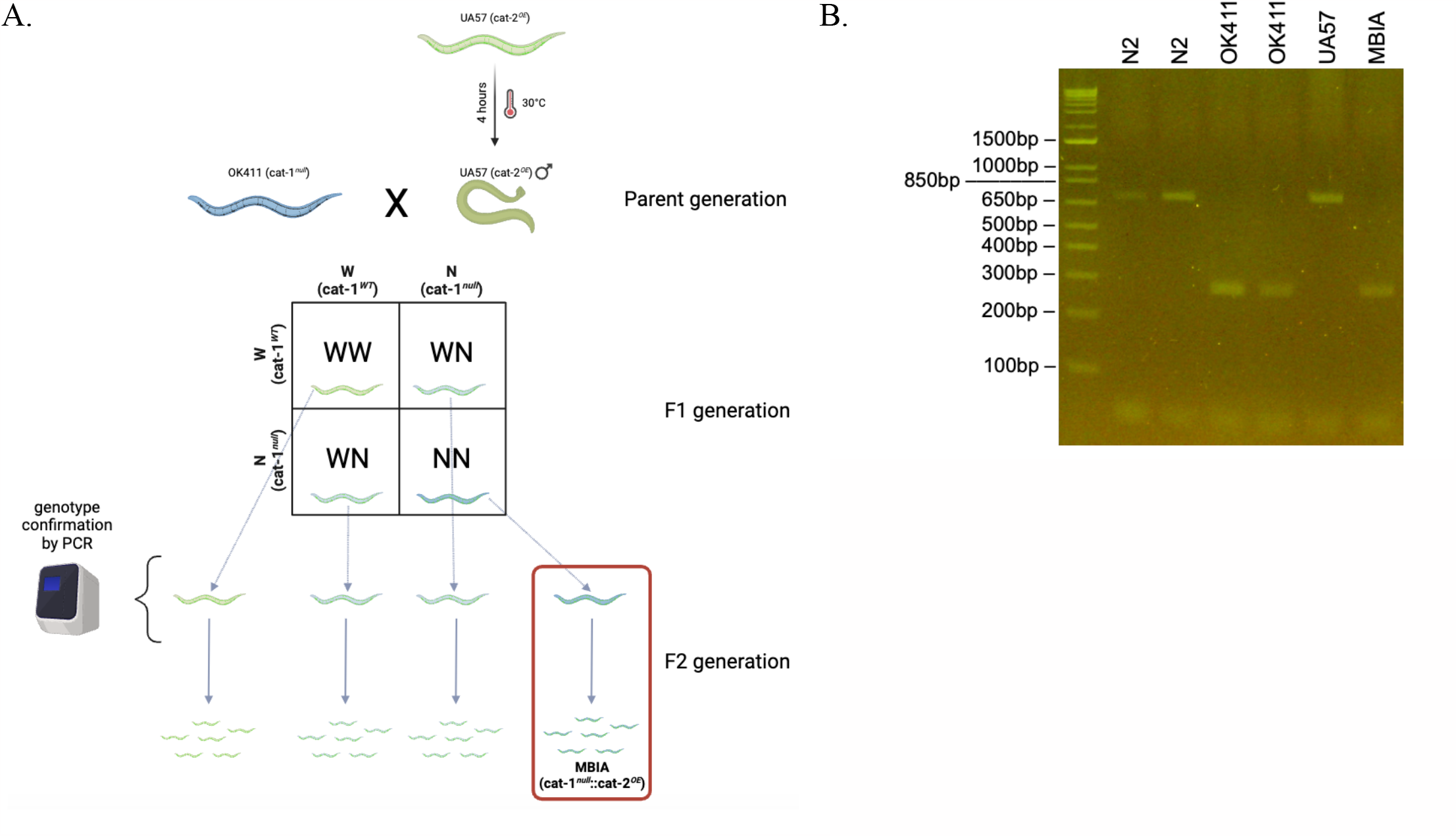
Creation of novel *C. elegans* strain “MBIA” overexpressing *cat-2* with *cat-1* null. A. Schematic demonstrating the process of generating MBIA (*cat-1*^*null*^*::cat-2*^*OE*^) strain from parent UA57 (*cat-1*^*WT*^*::cat-2*^*OE*^) and OK411 (*cat-1*^*null*^*::cat-2*^*WT*^) strains. Generation process consisted of heat-shocking UA57 C. elegans to generate males, which were bred with hermaphrodite OK411 C. elegans . Individual F1 generation C. elegans were selected and allowed to reproduce to generate single genotype lineages. B. PCR from single genotype lineage populations demonstrating *cat-1* gene product at 701 base pairs for N2 and UA57 strains. Gene deletion resulting in *cat-1* null phenotype results in 272 base pair gene product in OK411 and MBIA strains. Graphic made with BioRender.

**Figure 2.**
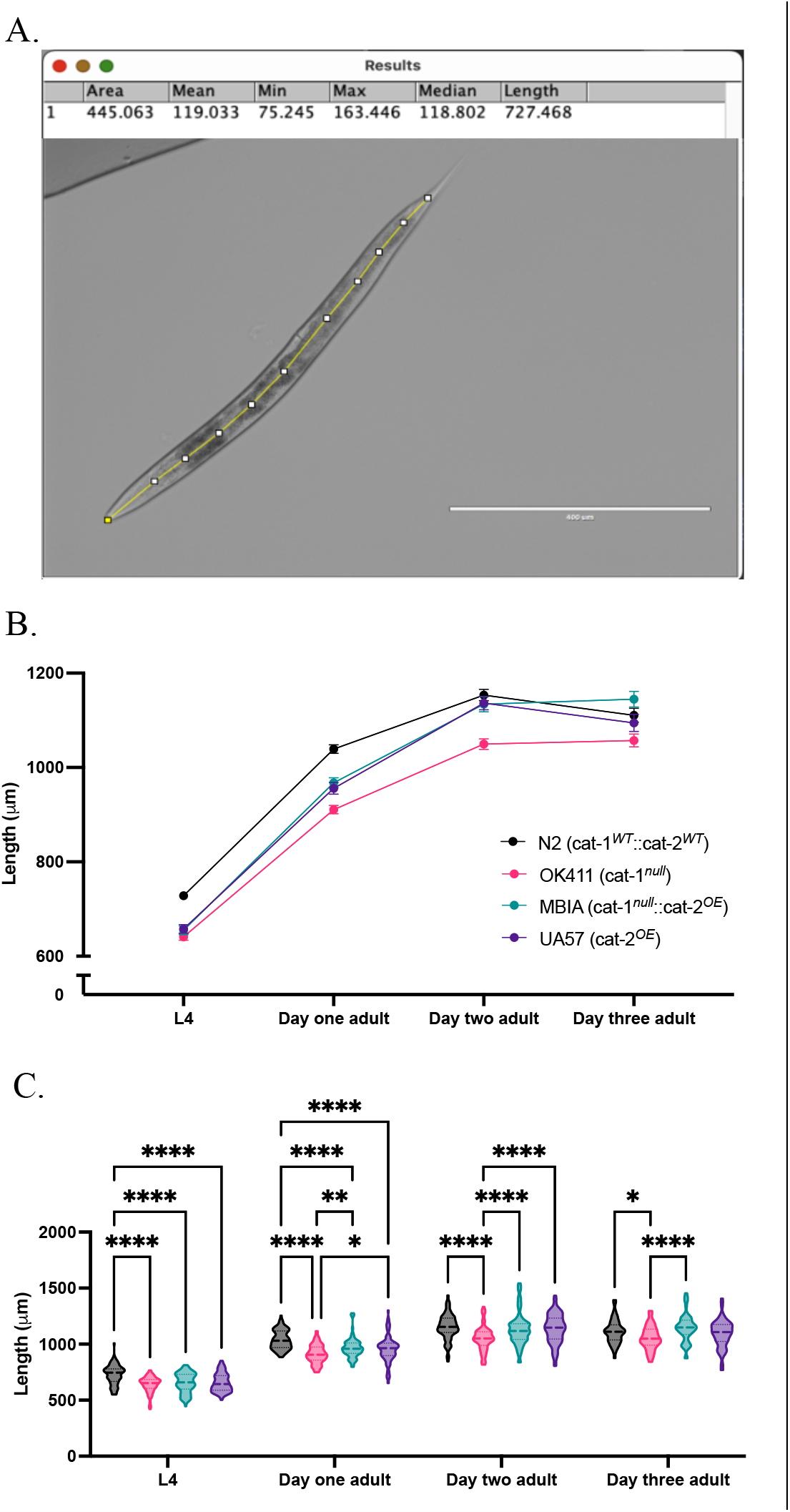
Dysregulation of dopamine synthesis and packaging effects *C. elegans* body length determined by microscopy. A. Representative brightfield image of *C. elegans* used for body length analysis in ImageJ with FIJI with segmented line through midline of animal to generate length measurement. B. Trend of body lengths for wild-type N2 (*cat-1*^*WT*^*::cat-2*^*WT*^) in black, OK411 (*cat-1*^*null*^*::cat-2*^*WT*^) in pink, MBIA (*cat-1*^*null*^*::cat-2*^*OE*^) in teal, UA57 (*cat-1*^*WT*^*::cat-2*^*OE*^) and in purple. tracked over four developmental stages. C. Violin plot with median and quartile demarcations demonstrating the body lengths for each strain at each developmental stage compiled across four experimental replicates with 47-93 C. elegans per genotype at each developmental stage for each experiment (total 1244 C. elegans analyzed). Two-way ANOVA with Tukey’s post-hoc test. Effect of developmental stage (F(9, 1228) = 4.209, p <0.0001); effect of genotype (F(9, 1228) = 1329, p <0.0001); effect of interaction (F(9, 1228) = 43.65, p <0.0001). *p <0.05, **p <0.01, ***p <0.001, ****p <0.0001.

### Dysregulation of dopamine synthesis and packaging affects C. elegans body length

To determine the effect of dopamine system mutations on body size, *C. elegans* populations were synchronized and allowed to develop to the L4 stage, day one adult, day two adult, and day three adult stages. At each stage, *C. elegans* were moved from agar culturing plate onto microscope slides with agar pads in a solution of Levamisole (5mM) to immobilize animals for brightfield imaging at 4x. Body length was determined by processing images in ImageJ with FIJI using the segmented line tool. Lines were drawn through the midline of each worm from head to tail, consistently beginning and ending at the same morphological points on each animal and reported in μm (Figure 1A). The body length of N2s were significantly larger than all other strains at the L4 stage (Two-way ANOVA with Tukey’s post-hoc test, n = 47-93 *C. elegans* per genotype at each developmental stage, p < 0.0001); however, there was no significant difference in length between the OK411, MBIA, and UA57 strains at the L4 stage. N2 nematodes were still significantly longer than OK411, MBIA, and UA57 *C. elegans* at the Day One Adult stage (Two-way ANOVA with Tukey’s post-hoc test, n = 47-93 *C. elegans* per genotype at each developmental stage, p < 0.0001), and OK411 nematodes were significantly shorter than MBIA (p<0.01) and UA57 (p<0.05) nematodes at the Day One Adult Stage. There was no significant difference in length between N2, MBIA, and UA57 strains at the Day Two Adult stage, and OK411 nematodes remained significantly shorter than the three other strains (p<0.0001). At the Day Three Adult stage, there was also no significant difference in length between N2, MBIA, and UA57 strains, and OK411 nematodes remained significantly shorter than N2 and MBIA strains (p<0.05 and p<0.0001 respectively).

As a secondary measure of body size, *C. elegans* were analyzed using the COPAS FP-250 large particle flow cytometer for high-throughput automated measurement. *C. elegans* were collected at L4 and Day One Adult stages for analysis of length. Body size was reported for each worm as time of flight (TOF) indicating length, and extinction (EXT) indicating width (Figure 3A). Following data processing detailed in methods section to remove any erroneous objects (e.g., contaminants), values were normalized within each experiment to N2 values to control for variations resulting from machine operational differences and outliers were removed. At the L4 stage, N2 and MBIA strains had significantly larger TOF values than OK411 and UA57 strains (p<0.0001); however, there was no significant difference in TOF between N2 and MBIA (Figure 3B) (One-way ANOVA with Tukey’s post-hoc test, n = 5789 total *C. elegans* compiled across six experimental replicates). At the Day One Adult stage, N2 and MBIA strains had significantly larger TOF values than OK411 and UA57 strains (p<0.0001); however, there was no significant difference in TOF between N2 and MBIA (Figure 3C) (One-way ANOVA with Tukey’s post-hoc test, n = 6899 total *C. elegans* compiled across six experimental replicates). At the L4 developmental stage, N2 had significantly larger EXT values than all other strains (p<0.0001), OK411 had significantly smaller EXT values than all other strains (p<0.0001), and UA57 had significantly larger EXT values than MBIAs (p<0.0001) (Figure 3D) (One-way ANOVA with Tukey’s post-hoc test, n = 5968 total *C. elegans* compiled across six experimental replicates). At the Day One Adult stage, OK411 EXT values remained significantly smaller than all other strains (p<0.0001), and N2 EXT values remained significantly larger than all other strains (p<0.0001).

**Figure 3.**
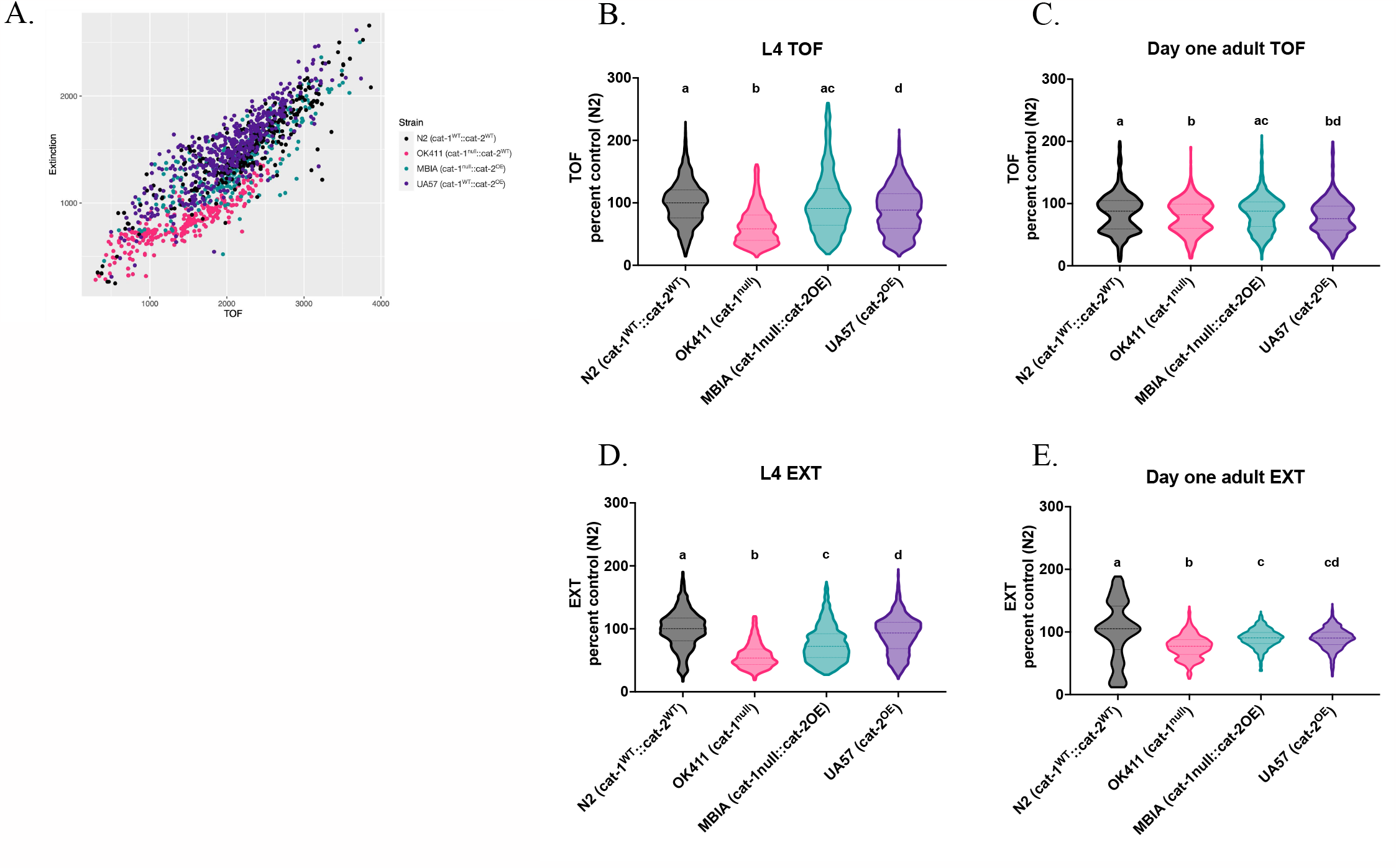
Dysregulation of dopamine synthesis and packaging effects *C. elegans* body length determined by automated analysis. A. Representative TOF vs. EXT scatter plot generated from cleaned COPAS data from L4 *C. elegans*. B. Normalized TOF values for L4 *C. elegans*. One-way ANOVA with Tukey’s post-hoc test n = 5789 total C. elegans compiled across six experimental replicates, F (3, 5785) = 76.38, a::b p <0.0001, a::d p <0.0001, b::c p <0.0001, b::d p <0.0001, c::d p <0.0001. C. Normalized TOF values for Day One Adult *C. elegans*. One-way ANOVA with Tukey’s post-hoc test n = 6899 total C. elegans compiled across six experimental replicates, F (3, 6895) = 15.57, a::b p <0.0001, a::d p <0.0001, b::c p <0.0001, c::d p <0.0001. D. Normalized EXT values for L4 *C. elegans*. One-way ANOVA with Tukey’s post-hoc test n = 5968 total C. elegans compiled across six experimental replicates, F (3, 5964) = 57.59, a::b p <0.0001, a::c p <0.0001, a::d p <0.0001, b::c p <0.0001, b::d p <0.0001, c::d p <0.0001. E. Normalized EXT values for Day One Adult *C*. with Tukey’s post-hoc test n = 3886 total C. elegans compiled across six experimental replicates, F (3, 3882) = 441.8, a::b p <0.0001, a::c p <0.0001, a::d p <0.0001, b::c p <0.0001, b::d p <0.0001.

There was no significant difference between MBIA and UA57 strains (p=0.9949) (Figure 3E) (One-way ANOVA with Tukey’s post-hoc test, n = 3866 total *C. elegans* compiled across six experimental replicates).

### Dysregulation of dopamine synthesis and packaging affects C. elegans vulval development and egg-laying

To distinguish whether the observed differences in body size had any effects on development or reproduction, synchronized populations of *C. elegans* were evaluated for vulval development and egg laying deficits. Vulval staging was performed at L4 stage with precise timing to ensure all strains were evaluated at the same time post-L1 plating. Vulval stages were observed using brightfield microscopy at 20x and staged based on morphological definitions according to previously established protocols.^36^ While the majority of N2 *C. elegans* had fully developed vulvas corresponding to stage 8 and 9, OK411, MBIA, and UA57 strains all were significantly delayed in vulval development (Figure 4A) (One-way ANOVA with Tukey’s post-hoc test, n = 11-20 *C. elegans* per genotype, p < 0.0001). It has been previously reported that *cat-1* null *C. elegans* have egg-laying deficits resulting in an increase in eggs *in utero*.^14,18^ To determine whether overexpression of *cat-2* could rescue egg-laying deficits observed in *cat-1* null animals, synchronized populations underwent a fecundity assay to quantify the number of eggs *in utero* at Day One Adult, Day Two Adult, and Day Three Adult (Figure 4B). At Day One Adult, UA57 had significantly fewer eggs *in utero* than N2 (p <0.0001), OK411 (p <0.05), and MBIA (p <0.0001), and OK411 had significantly fewer eggs *in utero* than MBIA (p <0.01) (Figure 4B) (Two-way ANOVA with Tukey’s post-hoc test, n = 45-61 *C. elegans* per genotype at each developmental stage). At Day Two Adult, MBIA had significantly more eggs *in utero* than N2 (p <0.01), OK411 (p <0.05), and UA57 (p <0.0001) (Two-way ANOVA with Tukey’s post-hoc test, n = 45-61 *C. elegans* per genotype at each developmental stage). By Day Three Adult, the only significant difference in eggs *in utero* was between N2 and UA57 with significantly fewer eggs *in utero* in UA57 compared to N2 (p <0.05) (Two-way ANOVA with Tukey’s post-hoc test, n = 45-61 *C. elegans* per genotype at each developmental stage).

**Figure 4.**
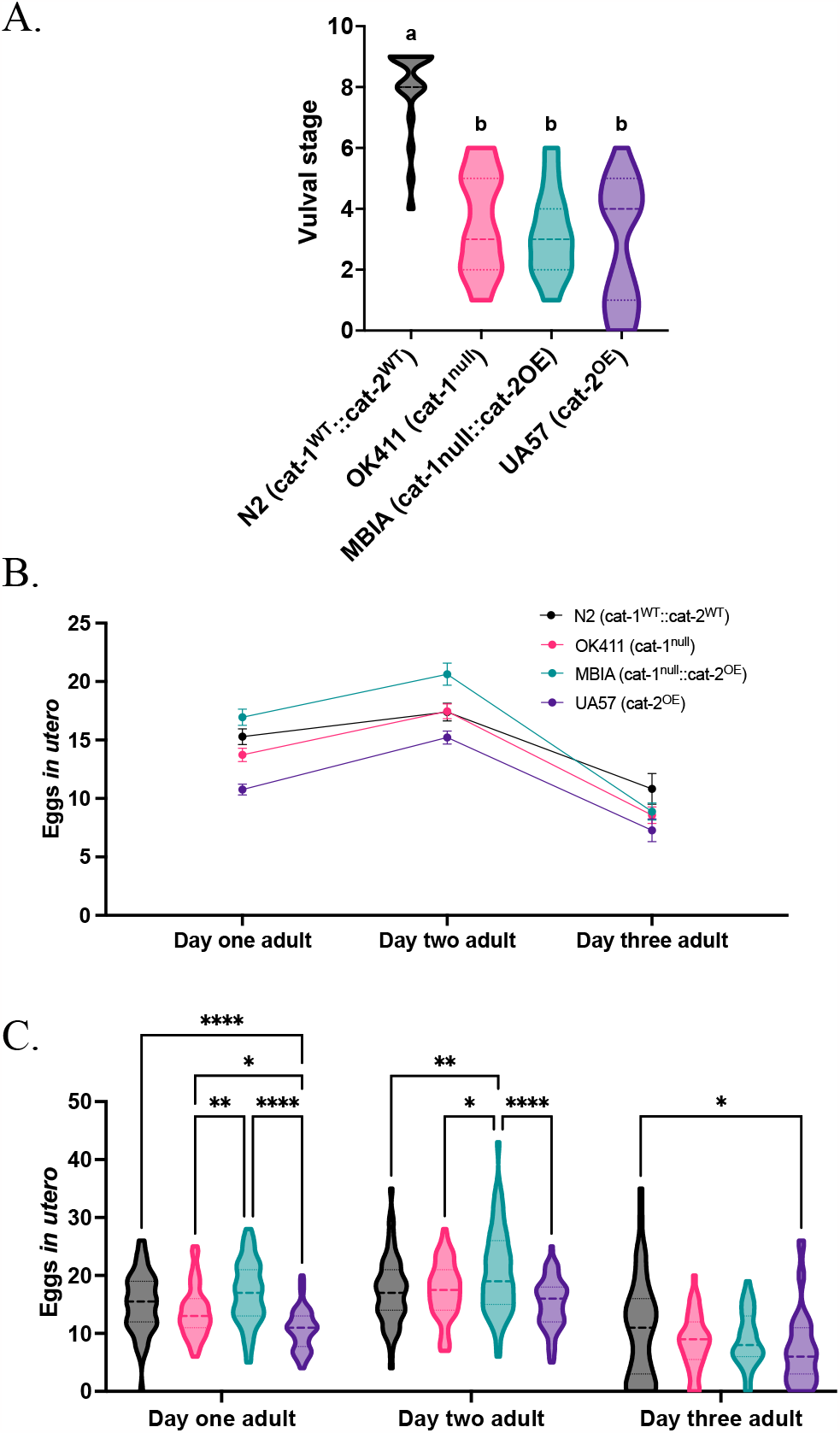
Dysregulation of dopamine synthesis and packaging affects *C. elegans* vulval development and egg-laying. A. Vulval stage determined by morphological analysis of L4 *C. elegans*. One-way ANOVA with Tukey’s post-hoc test n = 11-20 C. elegans per genotype compiled across two experimental replicates, F (3, 66) = 1.731, a::b p <0.0001. B. Trend of eggs *in utero* for wild-type N2 (*cat-1*^*WT*^*::cat-2*^*WT*^) in black, OK411 (*cat-1*^*null*^*::cat-2*^*WT*^) in pink, MBIA (*cat-1*^*null*^*::cat-2*^*OE*^) in teal, UA57 (*cat-1*^*WT*^*::cat-2*^*OE*^) and in purple. tracked over three developmental stages. C. Violin plot with median and quartile demarcations demonstrating the eggs *in utero* for each genotype at each developmental stage compiled across four experimental replicates with 45-61 C. elegans per genotype at each developmental stage for each experiment (total 650 C. elegans analyzed). Two-way ANOVA with Tukey’s post-hoc test. Effect of developmental stage (F(2, 6388) = 126.0, p <0.0001); effect of genotype (F(3, 638) = 18.41, p <0.0001); effect of interaction (F(6, 638) = 2.535, p <0.05). *p <0.05, **p <0.01, ***p <0.001, ****p <0.0001.

### Overproduction of dopamine partially rescues deficits in dopamine-mediated behaviors in C. elegans with loss of vesicular dopamine sequestration

As many behaviors in *C. elegans* have been previously demonstrated as mediated by dopamine, the effect of dopamine related genetic disruptions on *C. elegans* behavior was measured by quantifying curling and thrashing in 30s videos of swimming behavior performed at four developmental stages. Curling was defined as the complete overlap of a worm’s tail to a part of its body and quantified as number of curls in 30 sec intervals (Figure 5A).^35^ At L4, both OK411 and MBIA *C. elegans* demonstrated increased curling behavior compared to N2 (OK411 p <0.05, MBIA p <0.0005), with MBIA *C. elegans* also displaying increased curling compared to UA57 (p <0.01) (Figure 5B) (Two-way ANOVA with Tukey’s multiple comparisons test, n = 955 total *C. elegans* compiled across 3 experimental replicates). There were no significant differences in curling at the Day One Adult stage, and at Day Two Adult, OK411 had significantly more curling than N2 (p <0.01). At the Day Three Adult stage, UA57 demonstrated significantly more curling behavior than MBIA (p <0.05). Thrashing was defined as increased spasm-like behavior based on side-to-side movement during swimming behavior and was recorded as the number of seconds spent thrashing during a 30 sec interval (Figure 5C).^35^ At L4, OK411 spent significantly more time thrashing than N2 (p <0.005), MBIA (p <0.01), and UA57 (p <0.0001) (Figure 5D) (Two-way ANOVA with Tukey’s multiple comparisons test, n = 884 total *C. elegans* compiled across 3 experimental replicates). At L4, UA57 spent significantly less time thrashing than N2 (p <0.0001) and MBIA (p <0.0001) (Figure 5D). At the Day One Adult stage, OK411 continued to display significantly more time spent thrashing compared to N2 (p <0.0001), MBIA (p <0.0001), and UA57 (p <0.0001), and UA57 spent significantly less time thrashing than N2 (p <0.0001) and MBIA (p <0.0001). At Day Three Adult, the only significant difference between genotype was significantly more time spent thrashing in the OK411 compared to the UA57 (p <0.0001), and by Day Three Adult, UA57 spent less time thrashing than N2 (p <0.0001), OK411 (p <0.0001), and MBIA (p <0.0001).

**Figure 5.**
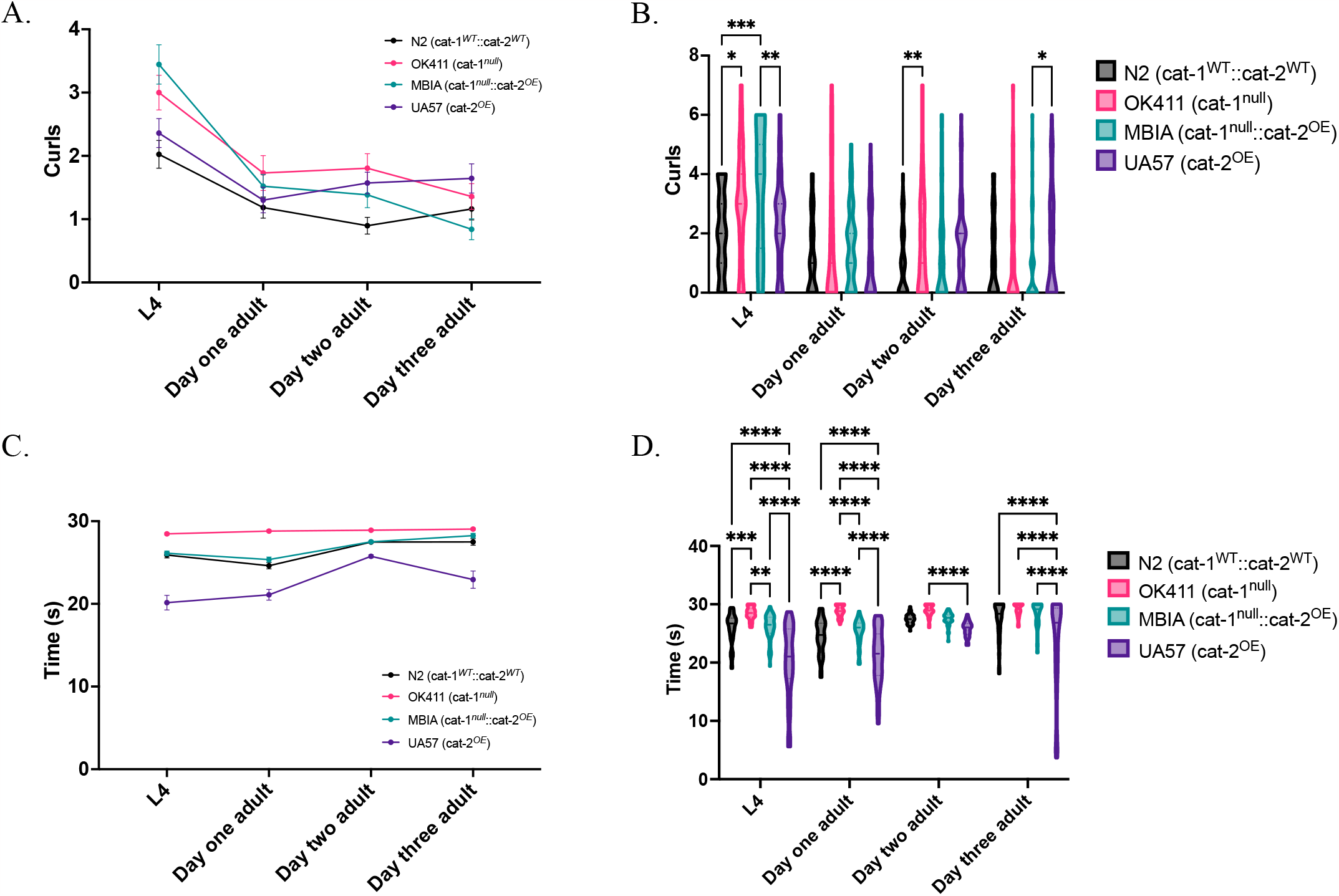
Overproduction of dopamine partially rescues deficits in dopamine-mediated behaviors in *C. elegans* with loss of vesicular dopamine sequestration. A. Trend of curls recorded in 30s interval of swimming behavior at four developmental stages with wild-type N2 (*cat-1*^*WT*^*::cat-2*^*WT*^) in black, OK411 (*cat-1*^*null*^*::cat-2*^*WT*^) in pink, MBIA (*cat-1*^*null*^*::cat-2*^*OE*^) in teal, UA57 (*cat-1*^*WT*^*::cat-2*^*OE*^) and in purple tracked over four developmental stages. B. Violin plot with median and quartile demarcations demonstrating the number of curls for each genotype at each developmental stage compiled across three experimental replicates with 955 total C. elegans analyzed. Two-way ANOVA with Tukey’s post-hoc test. Effect of developmental stage (F(3, 939) = 34.70, p <0.0001); effect of genotype (F(3, 939) = 6.768, p <0.0005); effect of interaction (F(9, 939) = 2.858, p <0.0001). C. Trend of time spent thrashing recorded in 30s interval of swimming behavior at four developmental stages. D. Violin plot with median and quartile demarcations demonstrating the time spent thrashing for each genotype at each developmental stage compiled across three experimental replicates with 884 total C. elegans analyzed. Two-way ANOVA with Tukey’s post-hoc test. Effect of developmental stage (F(3, 868) = 28.40, p <0.0001); effect of genotype (F(3, 868) = 131.0, p <0.0001); effect of interaction (F(9, 868) = 5.518, p <0.0001). *p <0.05, **p <0.01, ***p <0.001, ****p <0.0001.

### Overproduction of dopamine does not affect neuron size in C. elegans with loss of vesicular dopamine sequestration in early adulthood

Behavioral deficits resulting from dopamine dysfunction may be due to deficient dopaminergic neurotransmission or the degeneration of dopaminergic neurons. To determine whether the observed behavioral deficits resulted from differences in neuronal health, the size of dopaminergic neurons were quantified using GFP images of UA57 and MBIA *C. elegans* from the L4 to Day Three Adult stages. Neuron size was determined by processing GFP images in ImageJ with FIJI using the thresholding and ROI area tools. Thresholding was performed to create a mask over dopaminergic neurons and the area of the ROI was quantified (Figure 6A). There were no significant differences in neuronal size observed at any of the four stages (Figure 6B, C) (Two-way ANOVA with Šídák’s multiple comparison test, n = 1038 neurons measured across experimental replicates).

**Figure 6.**
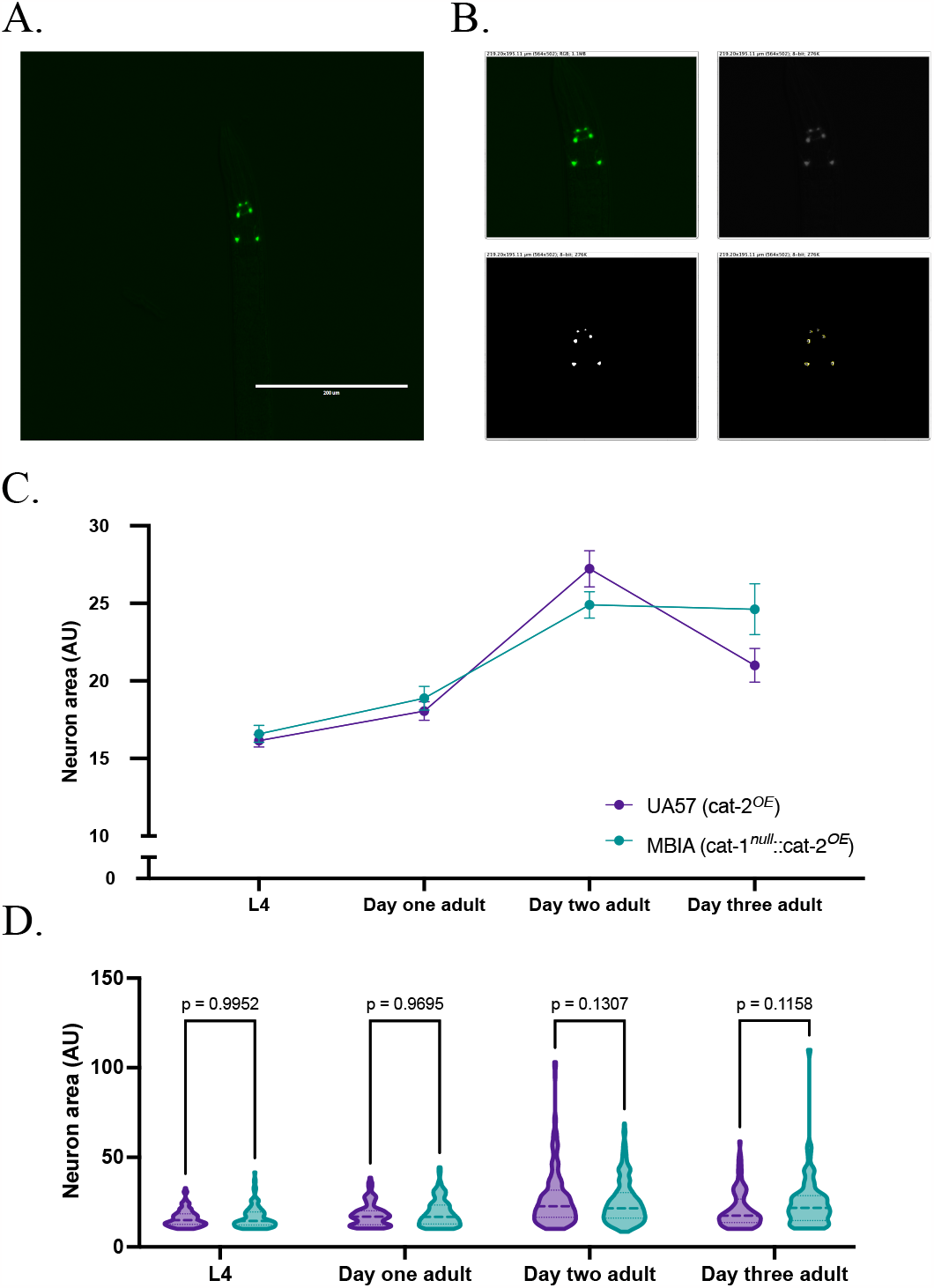
Overproduction of dopamine does not affect neuron size in *C. elegans* with loss of vesicular dopamine sequestration in early life. A. Representative 20x GFP image of dopaminergic neurons in the head of *C. elegans* used for neuron size analysis in ImageJ with FIJI. B. Image processing of original image (top left) demonstrating conversion to 8-bit image (top right), thresholding (bottom left), and ROI size analysis (bottom right). C. Trend of neuron area across four developmental stages with MBIA (*cat-1*^*null*^*::cat-2*^*OE*^) in teal, and UA57 (*cat-1*^*WT*^*::cat-2*^*OE*^) and in purple. D. Violin plot with median and quartile demarcations demonstrating the neuron size for each genotype at each developmental stage compiled across four experimental replicates. Two-way ANOVA with Tukey’s post-hoc test, n = 1038 total neurons measured across four experimental replicates. Effect of developmental stage (F(3, 1030) = 49.36, p <0.0001); effect of genotype (F(3, 1030) = 0.8204, n.s.); effect of interaction (F(3, 1030) = 3.202, p <0.05).

## Discussion

Building on previous publications that implicated dopaminergic signaling in the regulation of body size and behavior in *C. elegans*, we set out to determine whether increasing the endogenous production of dopamine could rescue deficits resulting from a lack of vesicular dopamine sequestration in *cat-1* null *C. elegans*. Prior studies discovered that *C. elegans* with *cat-2* null genotype, thus lacking the ability to catalyze the production of L-DOPA, had increased body size compared to wild-type animals.^37^ Interestingly, despite this study demonstrating an increase in body size due to lack of dopamine synthesis, we have anecdotally observed smaller body sizes in *C. elegans* with *cat-1* null phenotype that have normal dopamine production but lack the protein responsible for vesicular sequestration of dopamine. Here, we performed thorough analysis of body size by quantifying body length through two independent measures. Our data demonstrate that OK411 *C. elegans* lacking the *cat-1* protein have significantly shorter body lengths than wild-type N2 *C. elegans* throughout development and persisting into fully developed adulthood (Figures 2 and 3). *C. elegans* that overproduce dopamine through overexpression of the *cat-2* gene likewise showed shorter body lengths than wild-type at early adulthood stages (i.e., L4 and Day One Adult); however, their body length was not significantly different from wild-type *C. elegans* by Day Two and Day Three Adult stages (Figures 2 and 3). Thus, our data suggests that endogenous overproduction of dopamine is sufficient to rescue body size deficits resulting from deficient vesicular sequestration of dopamine.

To determine whether the observed differences in body length were a result of developmental delay, we investigated two measures of *C. elegans* development and reproduction: vulval staging and egg laying. Analysis of vulval staging performed at 48h post-L1 plating demonstrated a trend toward more fully developed vulva in N2 compared to fewer fully developed vulva in OK411, MBIA, and UA57 (Figure 4), suggesting a slight delay in the development of the vulva due to mutations in dopamine related proteins. However, by Day One Adult, all strains had begun to lay eggs (data not shown), suggesting that any delay in vulval development did not impair the laying of eggs by the Day One Adult stage. While previous studies have reported potential egg-laying deficits in *cat-1* null *C. elegans*, this was not observed in our analysis (Figure 4). However, analysis of eggs *in utero* did demonstrate significantly fewer eggs *in utero* in UA57 *C. elegans* at the Day One Adult stage, suggesting a potential enhancement in egg laying due to excessive dopamine production (Figure 4). By the Day Two Adult stage, MBIA *C. elegans* had significantly more eggs *in utero* compared to all other strains, which may suggest an egg-laying deficit; however, by the Day Three Adult stage, MBIA had no significant differences in eggs *in utero* compared to any other strains (Figure 4). While UA57 had no significant differences in eggs *in utero* compared to N2 at Day Two Adult, there were significantly fewer eggs *in utero* in UA57 at Day Three Adult, suggesting that fewer eggs *in utero* is a persistent phenomenon in UA57 *C. elegans* (Figure 4).

The data from automated high-throughput analysis performed using the COPAS biosorter confirmed the data obtained from manual analysis of microscopy images. Microscopy analysis was performed on *C. elegans* through the Day Three Adult stage by transferring synchronized adult *C. elegans* to new agar plates during each day of adulthood after egg-laying began to ensure no future generations of *C. elegans* were included in the analysis. Based on the method of collecting *C. elegans* and number of animals analyzed using the COPAS, analysis was only performed at L4 and Day One Adult stages to prevent the inclusion of future generations of *C. elegans* in the analysis. The COPAS biosorter typically uses the distribution of TOF and EXT to define and sort *C. elegans* based on developmental stages.^38^ Given that our microscopy data demonstrates that OK411 *C. elegans* show a persistent smaller body length throughout all developmental stages analyzed, it is important for experimenters to ensure there are no body size deficits resulting from genetic variation or pharmacological or toxicological treatment in experiments that utilize the COPAS biosorter for analysis or sorting of animals. We have reported that gravid adult OK411 *C. elegans* have a smaller body size than wild-type, demonstrating that body size cannot always be used as a proxy for developmental stage.

To understand the impact of gene variants in dopamine-related proteins, two additional behaviors (e.g., curling and thrashing during swimming) were analyzed as measures of dopamine-mediated behavior. While MBIA demonstrated an increase in curling behavior in early development (i.e., L4 stage), by the Day One Adult stage, there were no differences in curling behavior in MBIA compared to N2 despite a trend toward increased curling behavior in OK411 across multiple developmental stages (Figure 5). This suggests that the endogenous overproduction of dopamine resulting from *cat-2* overexpression can rescue the effects of *cat-1* null in *C. elegans*. In analysis of thrashing behavior, there was a consistent trend of more amount of time spent thrashing in OK411 animals across multiple developmental stages (Figure 5). In comparison, UA57 showed less time spent thrashing compared to N2 at L4, the Day One Adult stage, and the Day Three Adult stage (Figure 5). Taken together, the data from OK411 and UA57 provide evidence of dopamine-mediated thrashing behavior where less dopamine transmission resulting from *cat-1* null results in increased thrashing, whereas increased dopamine transmission resulting from *cat-2* overexpression causes less thrashing. In MBIA animals, thrashing is significantly less than OK411, but significantly more than UA57, and not different from N2 animals at L4 and the Day One Adult stage, suggesting that endogenously overproducing dopamine can rescue the *cat-1* null effect (Figure 5). At the Day Two Adult stage and Day Three Adult stage, MBIA are not significantly different from N2, further supporting the hypothesis that *cat-2* overexpression rescues *cat-1* null deficits (Figure 5).

It is known that dysregulated dopamine can be neurotoxic and lead to the degeneration of dopaminergic neurons in rodent and *C. elegans* models.^18,24,25^ While most of these models require aging of animals to observe neurodegeneration, neuronal health analysis was performed to determine whether the observed behavioral effects were a result of neurodegeneration. Both the overproduction of dopamine and the lack of dopamine sequestration independently result in increased cytosolic dopamine; thus, it is possible that the MBIA animals would have exacerbated neuronal vulnerability due to the compounded increase in cytosolic dopamine resulting from both *cat-1* null and *cat-2* overexpression. Analysis was performed only on the UA57 and MBIA animals as these animals express GFP in their dopaminergic neurons, allowing for quantification of neuron size, and the N2 and OK411 animals do not express GFP in their dopaminergic neurons. As there were no observed differences in neuron area between the two strains, it is likely that the observed differences in dopamine-mediated behaviors were solely the result of dysregulated dopamine synthesis and packaging (Figure 6). Although no differences in neuron area were observed at these developmental stages, it is possible that analysis performed at later ages would display enhanced neuronal vulnerability in the MBIA. In conclusion, here we present data from the analysis of *C. elegans* with various genetic mutations in dopamine-related proteins. We first demonstrated that apparent body length abnormalities resulting from deficient vesicular sequestration of dopamine can be rescued, as shown by the body length similarities between the MBIA strain and wild-type. We have also demonstrated that body size can be affected independently of developmental stage, which is an important consideration in the design of experiments utilizing *C. elegans* when a particular developmental stage is required. Lastly, we demonstrated that behavioral abnormalities resulting from disruptions in dopamine signaling can be rescued by enhancing dopamine production without compromising the health of dopaminergic neurons. Our results suggest a mechanism by which the excess production of dopamine in the cytosol of MBIA dopaminergic neurons can exit the neuron through non-evoked release mechanisms, thereby achieving synaptic transmission and action at post-synaptic receptors. For instance, previous work has shown a reversal of the plasmalemmal dopamine transporter (DAT), which typically facilitates presynaptic reuptake of dopamine from the synapse, but in cases of excess cytosolic dopamine, such as resulting from amphetamine treatment, can efflux dopamine from the cytosol into the synapse.^39^ The characterization of dopamine-related mutations of body size, development, and behavior can be used as a baseline to which *C. elegans* exposed to pharmacological and toxicological compounds can be compared to identify compounds that modulate dopamine signaling.

## Acknowledgments

We would like to acknowledge Joshua M. Bradner and Fion K. Lau for their assistance in worm husbandry and MBIA generation and Haejung Chung with her assistance with genotyping.

## Data availability statement

Data will be made available on Dryad upon publication.

